# Gum Arabic (*Acacia senegal*) enhances reproduction and modulates the microbiota-gut-brain axis of zebrafish in a sex-specific and dosage-dependent manner

**DOI:** 10.1101/2024.10.04.616708

**Authors:** Justin Abi Assaf, Jean-Charles de Coriolis, Alice May Godden, Eve Redhead, Jamie Bartram, Jayme Cohen-Krais, Karina Silova, Zoe Crighton, Gwenaelle Le Gall, Saber Sami, Sami Ahmed Khalid, Simone Immler

## Abstract

Dietary fibres (DFs) constitute a wide range of heterogeneous compounds that resist digestion and have beneficial effects on general health. Gum Arabic (GA) is a tree exudate consisting of 90% arabinogalactan, a polymer of arabinose and galactose sugars with prebiotic properties. As a dietary fibre, GA improves renal function, metabolism, and immune response in humans and animals. However, the underlying mechanisms leading to these health benefits are poorly understood. We supplemented female and male zebrafish (*Danio rerio*) with two concentrations of GA (6% and 60%) for two weeks. We assessed the effects of GA supplementation on the gut microbiome composition, intestinal and brain metabolic profiles, reproductive fitness, and brain gene expression. We found that GA supplementation resulted in changes to the gut microbiome with a relative increase in Fusobacteria and a relative decrease in Proteobacteria where the beneficial genus *Cetobacterium* was significantly more abundant after supplementation. GA supplementation increased acetate levels, particularly in the brain, causing a decreased expression of *cart1* in the brain of female zebrafish. While GA supplementation increased overall activity in male and female fish, reproductive fitness was negatively affected by GA supplementation in females. Our results suggest that while GA supplementation may have positive effects on metabolic rate and overall activity, it may come at a trade-off with reproductive fitness.

**Significance Statement:** Dietary fibres, found in plant-based food sources, can improve health. They include natural gums like gum Arabic, a highly sought-after food additive used as a homogeniser. Despite our better understanding of nutrition, a fibre gap is still prevalent in the Western world with efforts being made to incorporate new sources to close this gap and boost well-being. Here, we showed that when gum Arabic was supplemented into the zebrafish diet, it had a beneficial modulatory effect on the microbiota-gut-brain axis and reproductive fitness. Our findings support the benefits of dietary fibres but also link their impact to sexual dimorphism and dosage. This has implications for developing nutrition guidelines for both animals and humans.

## Introduction

A diet rich in dietary fibres (DFs) is thought to be essential for optimal health and well-being by preventing various non-communicable diseases due to their putative positive effects on cardiovascular, metabolic, and cognitive health in addition to weight management, appetite regulation, and inflammation (1–4). Gum Arabic (GA) is an edible tree gum exudate obtained by the incision of the trunks and the branches of *Acacia senegal* (L.) Willdenow native to the arid and semi-arid regions of Africa. Also known by several names like gum Acacia, Senegal gum, and Sudan Arabic gum, GA is chemically composed of highly branched and high molecular weight glycoproteins and polysaccharide complexes of which 90% is arabinogalactan, a polymer of arabinose (17-34%) and galactose (32-50%) with prebiotic properties, in addition to some amino acids and minerals (5, 6). It is a key additive (E-Number 414) in the food industry used as an emulsifier and a stabiliser, but its potential benefits as a DF have been largely overlooked. Traditional medicine has long suggested health benefits from its oral consumption, particularly with reports indicating positive effects on vital organ functions (7). Although not primarily consumed in the Western world as a source of DF, people in sub-Saharan regions like Sudan have incorporated GA into their day-to-day life for both medicinal and alimentary purposes long before it was considered safe for human consumption by regulatory bodies like the Food and Drug Administration (FDA) and the Food and Agriculture Organization (FAO) in the 1970s (8).

GA displays a range of potential health benefits. A randomised clinical trial (RCT) revealed that after 12 weeks of receiving 20 g/day of GA, the participants showed a significant decrease in systolic and diastolic blood pressure and fasting blood glucose (9). Similarly, the administration of GA was associated with improved renal function in patients with progressive chronic kidney disease (CKD) (7) and the oral administration of 30 g GA per day had antioxidant and anti-inflammatory effects in patients undergoing haemodialysis (10). A recently published systematic review of 29 clinical trials utilising GA as a therapeutic agent for various human diseases and conditions, indicated that GA improves oral health, and gastrointestinal functions, and supports healthy weight management in addition to alleviating adverse effects of sickle cell anaemia and rheumatoid arthritis (11). Beneficial effects have also been identified in mice where the administration of GA mitigated the expression of key pro-inflammatory plasma cytokines: IL-6, IL-1β, and TNF-α; despite being fed an inflammatory high-fat diet (12). Furthermore, GA improved semen quality in rats (13) and ovarian function in mice (14) while exhibiting no teratogenic effects (15); and in rats, GA enhanced cognitive ability whilst demonstrating neuroprotective properties (16, 17). Additionally, GA has prebiotic and modulatory effects on the gut microbiome (5, 18–20) via its fermentation by bacteria into microbial-derived metabolites, mainly short-chain fatty acids (SCFAs) (19, 21, 22). This microbiota-gut-brain axis is directly influenced by the diet which in turn can affect the cognitive behaviour and immunity of the organism (23–25). Given the broad range of health benefits linked to GA supplementation suggests that its effects likely occur at various levels between the gut-brain axis, but the exact mechanisms are not known.

In this study, we assessed the health implications of GA supplementation at multiple levels from the gut microbiome to brain function including the reproductive behaviour by experimentally manipulating the dietary supplementation of GA in zebrafish (*Danio rerio*). We exposed female and male adults (approximately 1 year old) to two GA-supplemented diets (6% or 60% GA) for two weeks and compared them to Control fish under standard diet. We assessed reproductive fitness and locomotory behaviour before collecting intestinal and brain samples for the analysis of the gut microbiome composition, tissue metabolic profiles, and brain transcriptome.

## Results

### The effects of GA on the gut microbiome

To assess the effects of GA on the microbiome and its potential prebiotic properties, we performed 16S sequencing of the gut microbiome. In line with previous findings, the dominant microbes in the zebrafish belong to two major phyla, Proteobacteria and Fusobacteria (26) (Fig. 1*A*). GA supplementation reduced the relative abundance of Proteobacteria in favour of an increase in Fusobacteria when compared to their respective controls and the strength of the effect was similar in 6% and 60% supplemented fish. However, the effects were more evident in females compared to males (Fig. 1*B*). The relative abundance of the main genus *Cetobacterium* in the Fusobacteria increased in females in both experiments whereas in males we only detected an increase in the 6% GA cohort (Fig. 1*C*). This genus is of particular interest as it is beneficial for the health of various teleost species including zebrafish where it has been shown to enhance immunity against pathogens, to reduce inflammation, and to produce SCFAs (27–30).

**Fig. 1.**
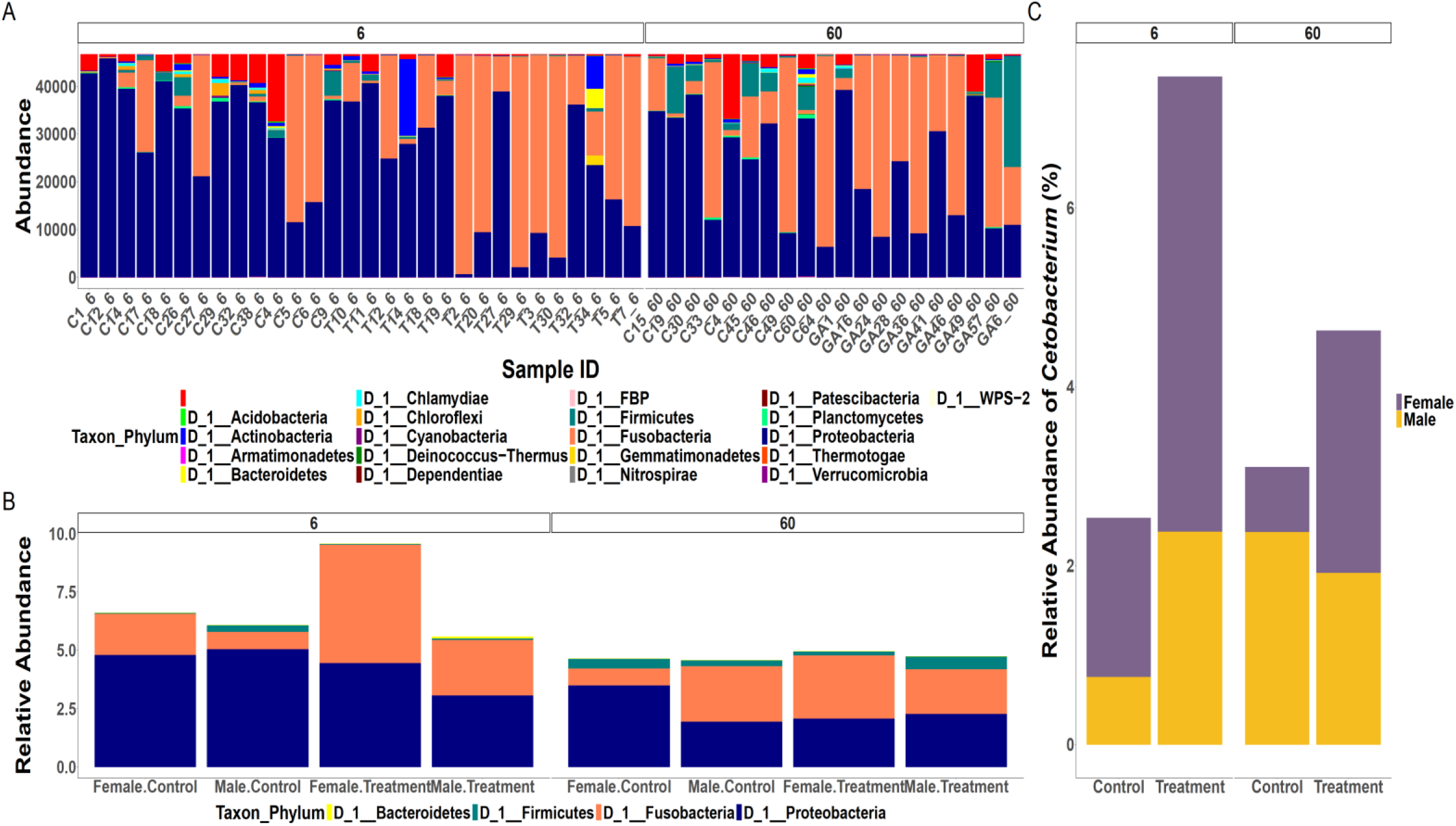
Microbiome composition stratified according to experimental condition and sex. (*A*) Abundance plot showcasing the two major phyla, Proteobacteria and Fusobacteria, in all zebrafish used in this analysis. (*B*) In both sexes, the Proteobacteria/Fusobacteria ratio drops in GA-supplemented fish compared to Control. (*C*) The increase in the relative abundance of *Cetobacterium* is more apparent in supplemented females when compared to their controls as opposed to the males.

We found no significant changes in α-diversity in either sex compared to their controls in the 6% GA experiment as indicated by the Observed Operational Taxonomic Unit (OTU) richness and the Shannon diversity index (Fig. 2*A* and *B*). However, the OTU index (but not the Shannon index) indicated a decrease in microbial diversity when comparing female fish from the 60% GA experiment to their control (Kruskal-Wallis, adjusted *p*-value = 0.003), but no such effect was apparent in males (Fig. 2*A* and *B*). This suggests that 60% GA can potentially reduce the number of gut bacteria in females without affecting their relative abundance. In 6% GA-supplemented fish, the unweighted and weighted UniFrac β-diversity were more uniform compared to Control fish as indicated by the tight clustering of samples and the narrow ellipses (Fig. 2*C* and *D*), and we found a significant effect of supplementation (PERMANOVA: F-value = 1.32, DF = 1, *p*-value = 0.043 and F-value = 3.3, DF = 1, *p*-value = 0.014) but no sex differences (PERMANOVA: F-value = 1.18, DF = 1, *p*-value = 0.1 and F-value = 2.24, DF = 1, *p*-value = 0.062), for unweighted and weighted UniFrac respectively. In fish under 60% GA supplementation, both females and males showed changes in the microbial communities compared to Control fish as indicated by the unweighted UniFrac β-diversity index (Fig. 2*C*), as a direct effect of the 60% GA dosage (PERMANOVA: F-value = 1.42, DF = 1, *p*-value = 0.021) but not sex (PERMANOVA: F-value = 1.02, DF = 1, *p*-value = 0.348). In contrast, the weighted UniFrac β-diversity index showed no significant differences between fish under 60% GA and Control (Fig. 2*D*; PERMANOVA for treatment: F-value = 1.09, DF = 1, *p*-value = 0.357 and PERMANOVA for sex: F-value = 1.60, DF = 1, *p*-value = 0.17).

**Fig. 2.**
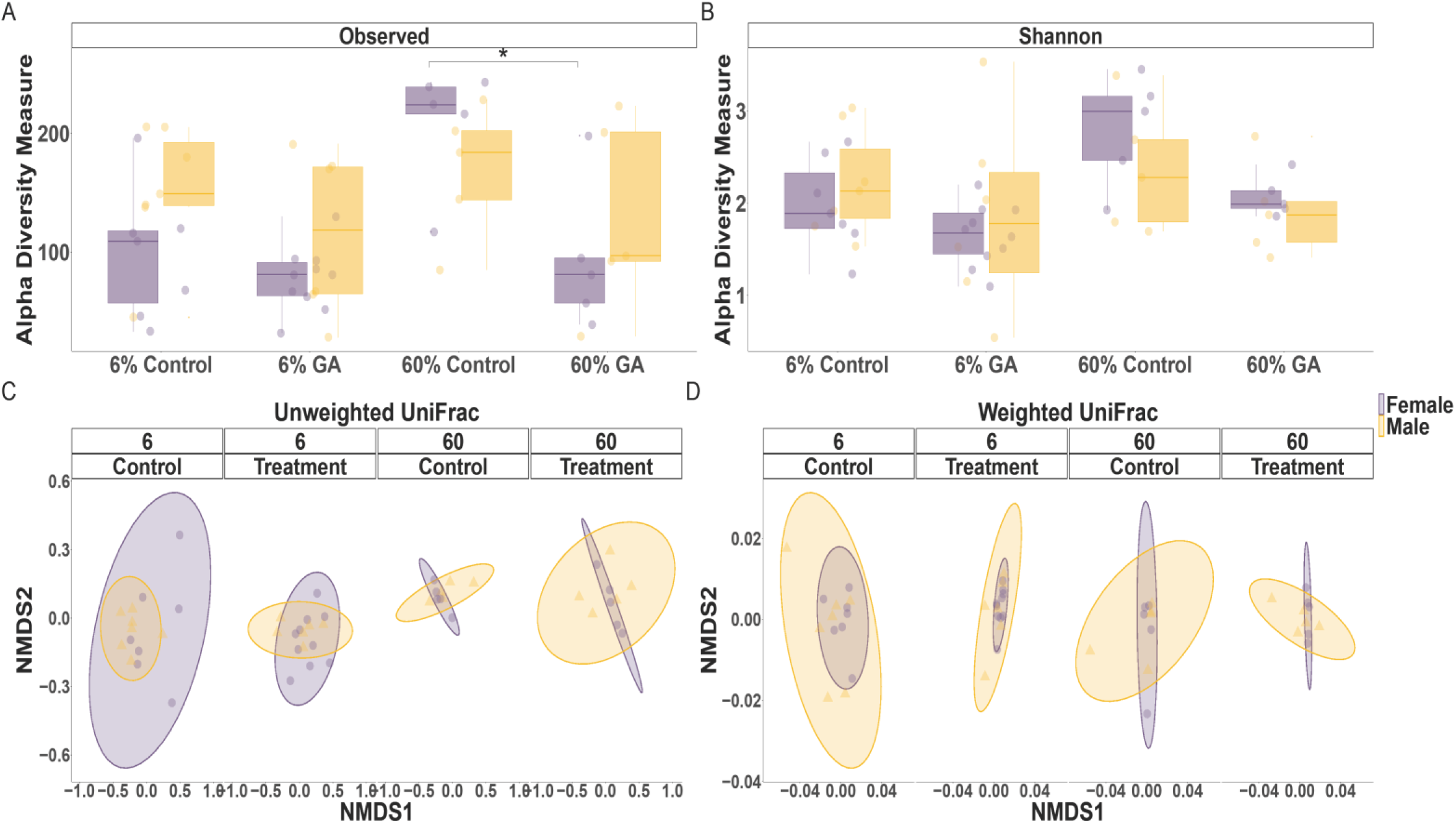
The α and β diversity are influenced by the GA supplementation, sex, and/or their interaction. (*A, B*) Observed and Shannon indices of gut microbiomes based on 16S analysis. Under 6% GA, there were no statistically significant changes in the α-diversity between the samples. Under 60% GA, only female fish displayed a significant drop (Kruskal-Wallis, * denotes adjusted *p*-value = 0.003) in α-diversity compared to the Control indicated by the Observed index. Each box plot represents the median, interquartile range, minimum, and maximum values with each dot representing a sample. (*C, D*) The unweighted and weighted UniFrac β-diversity based on 16S analysis. (*C*) Under 6% and 60% GA, the statistical difference in unweighted UniFrac was attributed to the GA supplementation (PERMANOVA, *p*-value < 0.05). (*D*) Under 6% GA, the statistical difference in weighted UniFrac was attributed to the GA supplementation (PERMANOVA, *p*-value < 0.05). However, under 60% GA, the weighted UniFrac was not changed by the GA supplementation, sex, or their interaction.

### The effects of GA on the metabolic profile of intestine and brain

We performed ^1^H NMR (nuclear magnetic resonance) metabolic analysis to test the effects of GA supplementation on the metabolome with a focus on the gut-brain axis. We identified and quantified 61 metabolites in the pooled samples (*SI Appendix*, Table S2 and Table S3) where 39 metabolites were detected in the intestinal samples and 22 metabolites were found in the brain (Fig. 3). Interestingly, acetate, one of the key SCFAs, was highly concentrated in the brain tissues of both female and male fish under 60% GA supplementation compared to Control fish. In the brain of GA-supplemented fish, we also observed a gradual increase in metabolites from the Control in both sexes with most of the cerebral metabolites being detected in the 60% GA fish. Interestingly, these detected brain and intestinal metabolites were most abundant in the 60% GA-supplemented female fish.

**Fig. 3.**
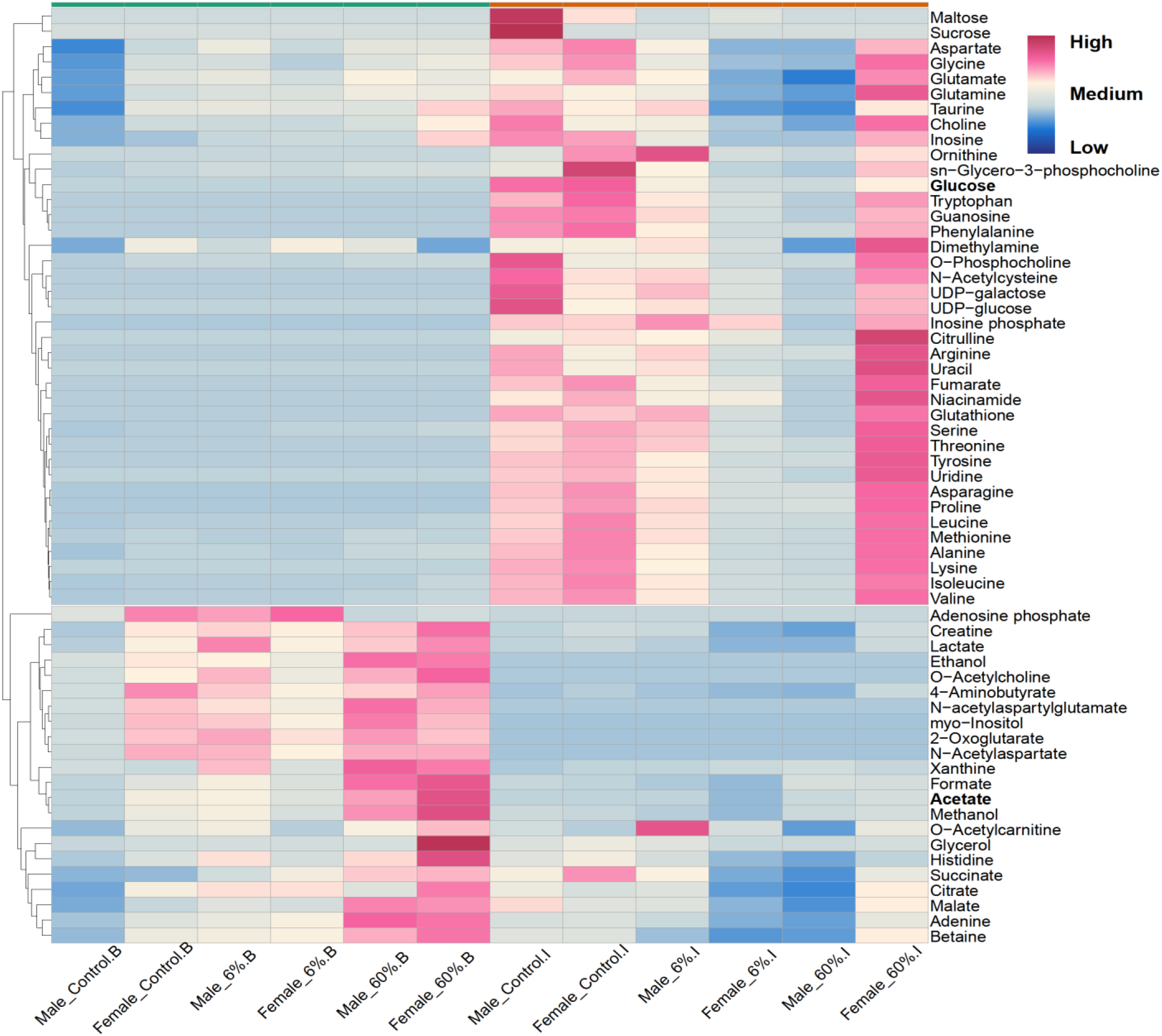
Heatmap displaying the detected 61 metabolites in the brain (B) and the intestine (I) by H^1^ NMR. The abundance of the metabolites shows a tissue-specific pattern. Colours in pink indicate a relative increase and colours in blue indicate a relative decrease.

### The effects of GA on organismal fitness

To test for the effects of GA supplementation on organismal fitness, we assessed reproductive fitness and general activity in the experimental adult fish and their offspring (Fig. 4*A-D*). At the end of the two-week dietary intervention, both Control and GA fish were crossed with non-experimental ABWT fish by natural spawning. Female zebrafish have a direct physiological contribution to clutch production by producing the eggs, whereas male zebrafish contribute indirectly by providing the sperm needed for egg fertilisation and by initiating the courtship and mating behaviour (31, 32). We found no statistically significant effects on clutch production (YES/NO) in fish under 6% GA supplementation (binomial Generalised Linear Mixed-Effects Models (GLMER): 0.36 [-0.2, 0.92], χ^2^ = 0.40, DF= 1, *p*-value = 0.52) nor 60% GA supplementation (binomial GLMER: -0.40 [-0.94, 0.114], χ^2^ = 0.54, DF = 1, *p*-value = 0.46) when compared to Control (Fig. 4*A*). Interestingly, the marginally non-significant interaction between 60% GA and sex may indicate a subtle increase in reproductive success in males compared to females (binomial GLMER: 1.32 [0.58, 2.06], χ^2^ = 3.18, DF = 1, *p*-value = 0.075) (Fig. 4*A*). Similarly, neither 6% GA (Poisson GLMER: -0.20 [-0.55, 0.15], χ^2^ = 0.10, DF = 1, *p*-value = 0.76) nor 60% GA supplementation (Poisson GLMER: -0.85 [-1.18, -0.52], χ^2^ = 0.13, DF = 1, *p*-value = 0.72) had an impact on the total number of eggs produced (size of clutch) compared to the Control group (Fig. 4*B*). However, under 60% GA, there was a significant interaction between treatment and sex (Poisson GLMER: 1.30 [1.24, 1.66], χ^2^ = 12.56, DF = 1, *p*-value < 0.001), indicating a sex-specific effect where GA supplemented females produced a significantly smaller clutch compared to 60% GA males (Pairwise difference of Treatment:Sex interaction using Poisson GLMER : -0.82 [-1.08, -0.557], DF = 1, *p*-value = 0.01) (Fig. 4*B*). Additionally, Control females produced larger clutches compared to the 60% GA-supplemented females (Pairwise difference of Treatment:Sex interaction using Poisson GLMER: 0.85 [0.52,1.19], DF = 1, *p*-value = 0.05) (Fig. 4*B*).

**Fig. 4.**
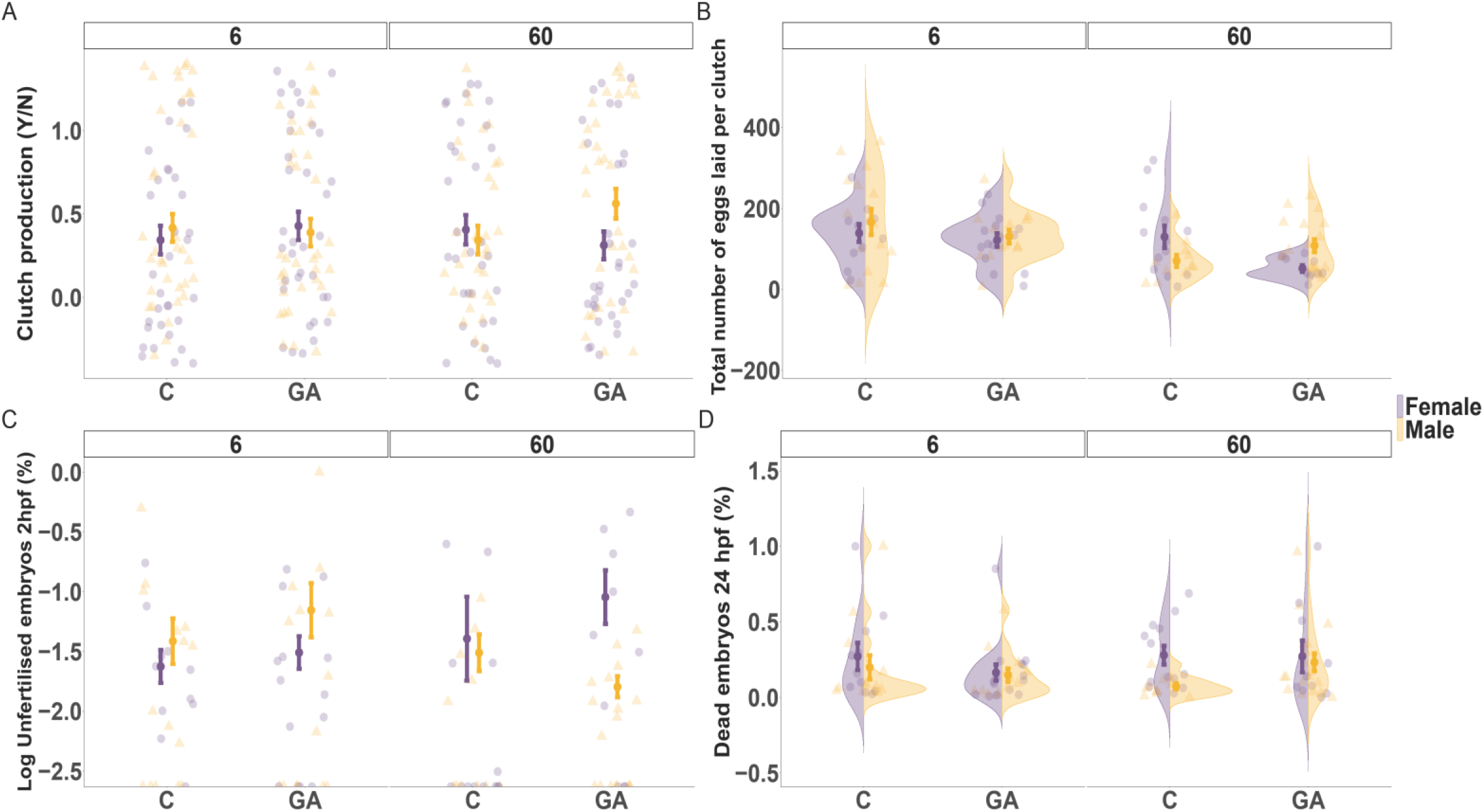
Dot plots and violin plots displaying the measures of reproductive fitness predominantly supporting male fitness at the expense of female fitness. (*A*-*D*) Each central point shows the mean value with the error bars representing the standard error of the mean (SEM). (*A*) Clutch production success during a spawning event with non-experimental fish. Compared to their controls, both 6% and 60% GA fish were unable to induce significant changes in clutch production. (*B*) No significant changes in the total number of eggs (clutch size) under 6% GA or 60% GA supplementation. A strong interaction between GA and sex was detected under 60% GA supplementation where supplemented females significantly produced fewer eggs per clutch compared to supplemented males. (*C*) Data are plotted on a logarithmic scale for better visualization with no change to the underlying data. The 6% GA supplementation produced no significant changes in the percentage of unfertilised embryos at 2 hpf. However, 60% GA females produced more unfertilised embryos compared to the Control chiefly attributed to the sex factor. Also, more unfertilised embryos were detected when comparing 60% GA females to the 60% GA males. (*D*) For embryo survival at 24 hpf, the effects were similar with no statistically significant changes between all comparison groups.

Whereas 6% GA supplementation had no significant effects on fertilisation success rate (binomial GLMER: -0.27 [-1.1, 0.56], χ^2^ = 0.08, DF = 1, *p*-value = 0.74) (Fig. 4*C*) or embryo survival at 24 hpf (hour post fertilisation) (binomial GLMER: -0.88 [-1.49,-0.27], χ^2^ = 0.55, DF = 1, *p*-value = 0.15) (Fig. 4*D*), 60% GA supplementation resulted in a higher ratio of unfertilised eggs in females but not in males compared to their Control mainly attributed to the sex effect (binomial GLMER: -0.14 [-0.88, 0.61], χ^2^ = 4.52, DF = 1, *p*-value = 0.03) rather than the Treatment:Sex interaction (binomial GLMER: -1.77 [-2.77, -0.77], χ^2^ = 2.92, DF = 1, *p*-value = 0.09) (Fig. 4*C*). Also, 60% GA females produced significantly more unfertilised embryos in comparison to the 60% GA males (Pairwise difference of Treatment:Sex interaction using binomial GLMER: 1.90 [0.96, 2.59], DF = 1, *p*-value = 0.03) (Fig. 4*C*). Surprisingly, despite the 60% GA males showing better fertilisation rates, embryo survival at 24 hpf was no different than the ones derived from the crossing of 60% GA females (Pairwise difference of Treatment:Sex interaction using binomial GLMER: -0.46 [-1.04, 0.12], DF = 1, *p*-value = 0.86) (Fig. 4*D*). Refer to Table S4 in *SI Appendix* for the full statistical models. Overall, it appears that whilst GA has potentially some beneficial effects on male reproductive fitness, the effects on female reproductive fitness, especially under high dosages, are detrimental.

When assessing the activity of the fish by measuring locomotion using artificial intelligence (AI) of video recordings of shoaling fish, we focused on 60% GA supplementation to associate this with the previously reported effects on reproductive fitness. We found that 60% GA-supplemented fish covered a significantly shorter total distance (mm) compared to Control fish (Gaussian Linear Mixed-Effects Models (LMER): -1281.20 [-1513.05, -1049.15], χ^2^ =30.49, DF = 1, *p*-value < 0.0001) (*SI Appendix*, Fig. S2*A*), but they swam faster (mm/s) (Gaussian LMER: 5.33 [3.98, 6.68], χ^2^ =15.53, DF = 1, *p*-value < 0.0001) (*SI Appendix*, Fig. S2*B*). This increase in activity level in the GA-supplemented fish could be a possible explanation for the contrasting effects on the reproductive fitness detected in females and males.

### The effects of GA on gene expression in the brain

We performed whole transcriptome analysis on brain tissues in female and male fish under 60% GA supplementation and tested for Differentially Expressed Genes (DEGs) (Table 1). When running a full model with treatment and sex as factors, we found no DEGs when comparing 60% GA to Control fish. However, the comparison of gene expression between Control females and Control males revealed three statistically significant DEGs (*ier2a, egr1*, and *egr4*), which were upregulated in females compared to males, indicating a difference in brain function, and supporting the sex-specific response to GA supplementation in the brains of females and males (33). Two genes, *cart1* and *slc1a2A*, were significantly downregulated in female fish when comparing 60% GA females with Control females but we found no DEGs when running the same analysis in males. When comparing gene expression in the brain between all fish under 60% GA supplementation and all Control fish without sex as a covariate, we found that *cart1* was once again downregulated whilst *snora74* and *clec3ba* were upregulated in the 60% GA fish compared to the Control fish. Interestingly, *cart1* has a role in feeding behaviour and appetite regulation (34). The detected downregulation of the *cart1* gene attributed to the sex effect, namely females in our case, further suggests a sex-specific brain regulation of hunger. Also, the positive ramification of GA on the brain was highlighted by the upregulation of *clec3ba* which is orthologous to human *CLEC3B* encoding for tetranectin, a protein with potential neuroprotective and immunomodulatory effects (35, 36).

**Table 1.** List of statistically significant differentially expressed genes identified in our analysis.

| Comparison | Gene name | Ensembl ID | Base mean | log2Fold Change | lfcSE (Standard Error) | adjusted p-value | Function |
| --- | --- | --- | --- | --- | --- | --- | --- |
| Control group only (female VS male) | ier2a | ENSDARG00000099195 | 460.0414 | 1.249918 | 0.232982 | $p < 0.01$ | Protein coding: Involved in several processes, including Kupffer's vesicle development; cilium assembly; and convergent extension. Localizes to nucleus. Is expressed in several structures, including axial chorda mesoderm; forebrain neural rod; midbrain hindbrain boundary; otic placode; and tail bud. Orthologous to human IER2 (immediate early response 2). (http://zfin.org/ZDB-GENE-030131-9126) |
| Control group only (female VS male) | egr1 | ENSDARG00000037421 | 4612.752 | 1.284819 | 0.24875 | $p < 0.01$ | Protein coding: Predicted to have several functions, including DNA-binding transcription factor activity, RNA polymerase II-specific; double-stranded DNA binding activity; and promoter-specific chromatin binding activity. Involved in embryonic retina morphogenesis in camera-type eye; embryonic viscerocranium morphogenesis; and glial cell development. Predicted to localize to cytoplasm and nucleus. Is expressed in head; heart; hindbrain neural keel; mesoderm; and nervous system. Human ortholog(s) of this gene implicated in asthma. Orthologous to human EGR1 (early growth response 1). (http://zfin.org/ZDB-GENE-980526-320) |
| Control group only (female VS male) | egr4 | ENSDARG00000077799 | 1441.57 | 1.483233 | 0.311542 | $p < 0.05$ | Protein coding: Predicted to have DNA-binding transcription factor activity, RNA polymerase II-specific and transcription regulatory region sequence-specific DNA binding activity. Predicted to be involved in positive regulation of transcription by RNA polymerase II. Predicted to localize to nucleus. Orthologous to human EGR4 (early growth response 4). (http://zfin.org/ZDB-GENE-080204-90) |
| 60 % GA VS Control (females only) | cart1 | ENSDARG00000070930 | 12.66215138 | - 5.655023705 | 1.137970828 | $p < 0.05$ | Protein coding: Predicted to have neuropeptide hormone activity. Predicted to be involved in several processes, including activation of MAPKK activity; |
|  |  |  |  |  |  |  | adult feeding behaviour; and negative regulation of appetite. Predicted to localize to extracellular space. Is expressed in brain; diencephalon; and hypothalamus. Human ortholog(s) of this gene implicated in obesity. Orthologous to human CARTPT (CART prepropeptide). (http://zfin.org/ZDB-GENE-030131-8025) |
| 60 % GA VS Control (females only) | slc1a2a | ENSDARG00000052138 | 9.292410708 | - 6.352755314 | 1.372234315 | $p < 0.05$ | Protein coding: Predicted to have symporter activity. Localizes to cone cell pedicle. Is expressed in brain; photoreceptor cell; and retinal neural layer. Human ortholog(s) of this gene implicated in early infantile epileptic encephalopathy 41. Orthologous to human SLC1A2 (solute carrier family 1 member 2). (http://zfin.org/ZDB-GENE-100422-11) |
| 60 % GA VS Control (all samples without sex as covariate) | SNORA74 | ENSDARG00000102120 | 0.463007 | 18.4799 | 2.941681 | $p < 0.001$ | Small nucleolar RNA |
| 60 % GA VS Control (all samples without sex as covariate) | cart1 | ENSDARG00000070930 | 7.480697 | - 4.71721 | 0.895641 | $p < 0.01$ | Protein coding: Predicted to have neuropeptide hormone activity. Predicted to be involved in several processes, including activation of MAPKK activity; adult feeding behaviour; and negative regulation of appetite. Predicted to localize to extracellular space. Is expressed in brain; diencephalon; and hypothalamus. Human ortholog(s) of this gene implicated in obesity. Orthologous to human CARTPT (CART prepropeptide). (http://zfin.org/ZDB-GENE-030131-8025) |
| 60 % GA VS Control (all samples without sex as covariate) | clec3ba | ENSDARG00000076541 | 118.9782 | 3.050037 | 0.648594 | $p < 0.05$ | Protein coding: Predicted to have carbohydrate binding activity. Predicted to be involved in bone mineralization. Predicted to localize to extracellular space. Human ortholog(s) of this gene implicated in osteoarthritis. Orthologous to human CLEC3B (C-type lectin domain family 3 member B). (https://zfin.org/ZDB-GENE-110411-71) |

## Discussion

Our results show that GA supplementation affects an organism at multiple levels, ranging from the gut microbiome to metabolic turnovers in the gut and the brain, ultimately changing overall fitness and behaviour. We found a dosage effect of GA, where a 6% GA supplementation had positive effects on the gut microbiome in both females and males with no implications on reproductive fitness, whereas 60% GA supplementation changed overall activity potentially at the cost of reduced reproductive fitness, particularly in females. This latter finding also suggests a sex-specific effect of GA supplementation. We discuss our findings in the context of the current push towards a fibre-rich diet as part of a healthy lifestyle and pinpoint some potential caveats.

The microbiota-gut-brain axis is increasingly recognised to play a key role in organismal health. Changes in the gut microbiota are associated with a range of neurological conditions such as Alzheimer’s, Parkinson’s disease, autism spectrum disorders (ASD), and potentially even mood (37, 38). GA has previously been shown to enhance cognitive abilities and more recently to be anti-epileptic in rats (39). Our results suggest that the substantial changes in gut microbiota in GA-supplemented fish after just two weeks may have downstream effects and ramifications from gut health to brain function and behaviour. Due to the highly dynamic nature of the gut microbiome, dietary changes and the availability of nutritional substrates influence its composition and architecture. DFs have been thoroughly documented to exhibit gut modulatory effects in humans and animals, which contribute to their positive health benefits (40, 41). In line with these previous findings on other DFs, we showed that GA can favourably shift the Proteobacteria/Fusobacteria (P/F) ratio towards the more beneficial Fusobacteria and, consequently, *Cetobacterium* (Fig. 1*A*-*C*). In zebrafish, a drop in Fusobacteria and a consequential increase in the P/F ratio is directly linked with gut dysbiosis (42). Even a dosage of 6% GA was sufficient to generate beneficial effects on the gut microbiome over just two weeks. Our findings suggest a possible difference in the gut microbiome composition according to sex but mainly highlight the sex-specific response to GA supplementation.

The role of sex in shaping the gut microbiome is contested and inconsistent (43–47). Yet, research has shown that sex-specific hormones can alter the gut microbiome composition by influencing the host-microbiome interactions. At the same time, the sex hormones differentially affect the physiology and behaviour of the two sexes, which in turn affects the microbiome (43, 48, 49). However, a variation in diet has been consistently shown to exert sex-specific changes on the gut microbiome across different species including the zebrafish (50–52). In line with the idea of a sex-specific microbiome, we showed that the increase in the relative abundance of *Cetobacterium* was more apparent in females in both GA treatments. However, the higher dose of 60% GA only reduced the Observed α-richness in females (Fig. 2*A*). Since the zebrafish display sexual dimorphism in anatomical, physiological, and behavioural aspects (53–55), we speculate this sex-specific effect of GA on the microbiome to be due to diet-host interactions. *Cetobacterium*, a prominent bacterium in the Fusobacteria phylum, is known to produce acetate (56) which can cross the blood-brain barrier to control appetite and glucose metabolism (56, 57).

Our metabolic profiling analysis showed abundant levels of intestinal tissue glucose in the Control group as opposed to negligible amounts in the GA-supplemented fish (Fig. 3). Interestingly, this suggests that GA can regulate glucose metabolism and appetite via the microbiota-gut-brain axis. We propose that the acetate generated by *Cetobacterium* in response to GA supplementation induced insulin secretion by stimulating the parasympathetic nervous system (56). Subsequently, insulin reduces blood glucose levels by stimulating the glycolytic activity of the fish gut; thus, using up intestinal glucose storage for metabolism (58). Additionally, acetate plays a role in appetite and feeding behaviour which subsequently regulates metabolism via interactions with neuropeptides and gut hormones; however, these contrasting effects are dependent on the species, route of administration, and acetate source (57, 59–61). Notably, high levels of acetate were detected in the brain tissues of both females and males under 60% GA supplementation (Fig. 3). This indicates that GA can support metabolic health and regulate appetite in both sexes.

Our results provide further evidence for a positive effect of soluble DFs, like GA, on the microbiota-gut-brain axis. The observed increase in the relative abundance of *Cetobacterium* under 60% GA in female zebrafish may be linked to the downregulation of the *cart1* gene, anorexigenic gene, in female brain tissues via an acetate-dependent pathway. The effects of acetate on the regulation of the *cart1* gene remain inconsistent with some studies indicating no significant change (62) whilst others show an up-regulation (61) in gene expression. This discrepancy emphasises that the acetate-mediated effects on metabolism and appetite can differ depending on whether the source is microbial-derived or exogenous (59). From a mechanistic perspective, our study suggests that the microbial-derived acetate from the GA fermentation can potentially counter the anorexigenic effects of the *cart1* gene particularly in females. Furthermore, DFs have been shown to exhibit immunomodulatory effects on the gut and to shape the homeostasis of the immune system (63, 64). We detected the upregulation of the gene *clec3ba* in 60% GA-supplemented fish compared to the Control fish (Table 1), which is in line with the idea that GA may have an immunomodulatory effect and increase the expression of C-type lectins in the innate immune cells namely macrophages located in the zebrafish brain (65, 66). Incorporating GA into the zebrafish diet can potentially activate defence mechanisms against infections.

Overall health is often directly linked with reproductive fitness (67, 68). We found potentially beneficial effects of GA supplementation on male reproductive fitness, particularly at high dosages, whereas the effects were detrimental in females. In animals, the implications of DFs on the reproductive parameters of both sexes have been promising. In boars, the incorporation of inulin and wheat bran during the pre-puberty period enhanced testosterone and semen production (69). Also, in pigs, the incorporation of a variety of DFs primarily before mating, enhanced reproductive fitness and offspring survival by improving oocyte maturation (70). The health implications of DFs on human reproduction are currently understudied as most research focuses on metabolic health, but a high-fibre diet can reduce the reabsorption of oestradiol in the gut and subsequently be detrimental to the menstrual cycle in women due to the diet’s influence on the hypothalamic-pituitary axis, supporting a potential role of the gut-brain axis in reproduction (71, 72). Moreover, female gonadal hormones can have contrasting effects; therefore, low intake of DF may negatively affect male fertility (73). A possible explanation for the positive impact of GA on male reproductive fitness in our study is that it improves sperm quality as previously reported in mammals (13, 74, 75). Also, GA can influence the steroid hormones governing gametogenesis (75–77) and exhibit antioxidant activity to protect the gametes from damage (14, 16, 74, 77) in both males and females. Despite potentially displaying some adverse effects under specific conditions, DFs play a crucial role in safeguarding metabolic health which in turn can have positive ramifications on fecundity and reproductive nutrition (78, 79). The association between DFs and reproduction is not linear and likely dosage-dependent which warrants further investigation. Our study highlights the importance of considering differences in female and male physiology when making dietary recommendations and the possible role the microbiota-gut-brain axis has on reproductive fitness (48, 80–82).

One possible reason for the associations detected between the 60% GA dosage and reproductive fitness may be the behavioural changes we identified. We observed that fish supplemented with 60% GA exhibited increased speed (mm/s) but without swimming a longer distance (mm) compared to Control fish (*SI Appendix*, Fig. S2*A* and *B)*. We propose two different possible explanations for the increased speed in GA-supplemented fish. The first theory is that this hyperactivity may benefit males during courtship and spawning activities, where the males rapidly chase females whilst nudging their flanks with their snout and attempting to lead them to the spawning site (83). The second theory relates to foraging and prey-capture behaviour (84, 85), which could be linked to the downregulation of the *cart1* gene (anti-anorexigenic effect) in the 60% GA-supplemented females and may induce an increased desire for food or a change in feeding behaviour (86–88) (Table 1). The increase in activity under GA supplementation could in turn explain the reduced reproduction caused by energy allocation favouring somatic growth over ovarian growth and reproduction (89–91).

A diet rich in adequate DFs from whole-food plant-based sources yields downstream health benefits on our gut microbiome, metabolic, and cognitive functions. Our results suggest that DFs such as GA, can positively affect the gut microbiome composition and subsequently influence the brain gene expression and functions. However, we also found negative effects on female reproductive fitness at very high dosages. It is important to understand when and how DFs exhibit optimal benefits and to improve our understanding of the crosstalk between nutrition and health. Our study took a first stab in this direction and demonstrated that a change in DF, even over a short period, has substantial effects on the microbiota-gut-brain axis and even reproduction, but in a dosage-specific manner. More in-depth studies are warranted to further verify our findings and to establish sex-specific guidelines for optimal fibre intake.

## Materials and Methods

### Animals and Ethics

ABWT zebrafish were obtained from the European Zebrafish Resource Center (EZRC) in Karlsruhe, Germany. The fish were maintained at a 1:1 sex ratio under standard conditions in compliance with the UK Home Office guidelines in the Controlled Environment Facility (CEF) at the School of Biological Sciences, University of East Anglia (UEA). Experiments were performed with approval from the UK Home Office under project licence: P0C37E901.

### Genomic DNA Extraction and 16S rRNA Sequencing

Genomic DNA was isolated from n = 50 adult zebrafish intestines (14 Control fish and 16 6% GA fish/ 10 Control fish and 10 60% GA fish) using FastDNA™ SPIN Kit for Soil (MP Biomedical, product code:116560200-CF) as per manufacturer’s recommendations (*SI Appendix*, Fig. S1). This genomic DNA was sent to Source BioScience (UK) in Cambridge (Source Bioscience Sequencing, William James House, Cowley Road, Cambridge, CB4 0WU) for 16S rRNA sequencing (*SI Appendix*, Table S1) with 250 or 300 base pair-end reads. The sequencing platform used was Illumina MiSeq. The bioinformatic microbiome analysis pipeline QIIME2 was used to attain expression data (92). The R package *phyloseq* 1.44.0 (93) was used for the downstream analysis.

### Metabolites Extraction and ^1^H NMR Metabolomic Analysis

Due to the small amount of tissue in each organ, we pooled brains and intestines from two or three fish per treatment and sex, resulting in six pooled samples per tissue (*SI Appendix*, Table S2). The extraction protocol was conducted and optimised in the Le Gall lab as previously described (94). High-resolution [^1^H] NMR spectra were recorded on a 600-MHz Bruker Avance spectrometer fitted with a 5-mm TCI proton-optimized triple resonance NMR inverse cryoprobe and a 24-slot autosampler (Bruker, Coventry, England). All identified metabolites (*SI Appendix*, Table S3) were quantified using Chenomx NMR suite 8.6 software.

### Phenotypic Characterisation and Behavioural Assessment (AI)

The 6% GA experiment included a total of n = 144 fish (72 Control and 72 GA), whilst the 60% GA experiment included a total of n = 128 fish (64 Control and 64 GA) of equal sex ratio (*SI Appendix*, Fig. S1).

At the end of the two-week experimental period, natural spawning was initiated by pairing each experimental fish with a non-experimental fish of the opposite sex. The fish were placed in external breeding tanks, separated by a divider, the day before spawning. The breeding tanks were also covered with an opaque cloth to ensure darkness until the next day. On the morning of the experiment, after removing the opaque cover and dividers, the breeding tanks were checked hourly for eggs. In the event of success, the eggs were collected from the tanks and incubated at 28 °C in a 0.01% solution of system water and methylene blue. The resulting clutches were assessed for the different fitness traits at two time points: 2 and 24 hpf.

In the 60% GA experiment, fish were recorded for 60 minutes in an isolated room whilst being placed in a special tank. All recordings were captured by a GoPro HERO4, and tracking was performed with the idtracker.ai software (95). The specific settings for the AI tracking are mentioned in the *SI Appendix*, Materials and Methods.

### RNA Extraction and RNA-seq

Brain samples from a total of n = 40 fish (20 Control fish and 20 60% GA fish) were collected by dissection following euthanasia and kept cool over ice before freezing in liquid nitrogen (*SI Appendix*, Fig. S1). RNA extraction was performed using Quick-RNA™ Miniprep Plus Kit with Zymo-Spin™ IIICG Columns (Capped) & Spin-Away™ Filters (Zymo Research, product code: R1057, Cambridge Bioscience, UK) according to manufacturer’s recommendations. Library construction was performed using a NEBNext Ultra II Direction RNA-seq library kit (poly-A selection, 200-300bp inserts) by the company Novogene (UK) in Cambridge (25 Science Park, Milton, Cambridge, CB4 0FW). Illumina sequencing PE150 was used, resulting in each sample having a minimum of 300 reads and 6G of raw data per sample. The bioinformatics pipeline “nf-core/rnaseq” version: 3.11.1 (https://nf-co.re/rnaseq/3.11.1) (96) was used to analyse the RNA sequencing data obtained from the zebrafish brain samples using the Ensembl reference genome file (Danio rerio.GRCz11.109). The DEG analysis was performed using the *DESeq2* package (97). The analysis was stratified into five comparisons according to the diet condition and/or sex by changing the design formula of the *DESeqDataSetFromMatrix* function.

### Statistical Analyses

R (version 4.3.1) in R Studio 2023.06.0 Build 421 was used for all the analyses. The Kruskal-Wallis test was used for the α-diversity and statistical significance was determined with an adjusted *p*-value < 0.01. Statistical significance in *β*-diversity was determined using PERMANOVA with a *p*-value < 0.05. GLMM and LMER models from the library *lme4* were used for the phenotypic and AI data, respectively (98). Greater detail regarding these analyses can be found in the *SI Appendix*, Table S3. Statistical significance for the differentially expressed genes was determined by a log2 fold change with an adjusted *p*-value < 0.05.

## Data Availability

The 16S rRNA and RNA-seq sequence data for this project are available on GEO: GSE245252 and GEO: GSE245568, respectively (https://www.ncbi.nlm.nih.gov/geo/). All data are included in the manuscript and/or supporting information. Detailed materials and methods are fully presented in the *SI Appendix*.

## Acknowledgements

We would like to express our sincere gratitude to Ms. Rispah N. Ng’ang’a for her assistance with the experimental work and data collection. We thank Dr Sarah Worsley and Mr Chuen Lee from Prof David Richardson’s group at UEA for their help with the 16S microbiome analysis. All the bioinformatics tools presented in this paper were carried out on the High-Performance Computing (HPC) Cluster supported by the Research and Specialist Computing Support service at UEA. We also would like to thank the Disease Modelling Unit (DMU) staff at UEA for their assistance with the zebrafish husbandry. This work was supported by our funders: the European Research Council (ERC), the Natural Environment Research Council (NERC), and the Wellcome Trust.

