## Supporting Information for "Gum Arabic (*Acacia senegal*) enhances reproduction and modulates the microbiota-gut-brain axis of zebrafish in a sex-specific and dosage-dependent manner"

**This PDF file includes:**

Supporting text (Methods and Materials)
Figures S1 to S2
Tables S1 to S4
SI References

#### Materials and Methods

**Zebrafish Husbandry.** Eight adult fish were housed per 3.5 L tanks (6% GA experiment) and 16 adult fish per 8 L tanks (60% GA experiment) on a recirculating rack system (ZebTEC rack with Active Blue technology, Tecniplast, UK) at 28 °C and kept on a 14:10 hour light: dark cycle. *Ad libitum* feeding of the fish occurred three (weekend) to four (weekdays) times per day with a mixture of standard dry feed (ZEBRAFEED 400-600 by SPAROS, Área Empresarial de Marim, Lote C, 8700-221 Olhão, Portugal) and live *Artemia* (Sep-Art Artemia Cysts, Ocean Nutrition).

**Gum Arabic.** The GA used in the present study was collected in Sudan and taxonomically identified as *Acacia senegal* var. *senegal*. The Gum Arabic Board of Sudan provided it in spray-dried powder format. GA was fed to the fish by supplementing their dry food. Briefly, the desired amount of GA was weighed and dissolved in a minute amount of distilled water, enough to allow the GA to be dissolved entirely. This mixture was supplemented with a weighed amount of standard feed at a specific percentage (6% or 60%), creating a paste that was then spread out to form thin sheets and left for drying. Using a coffee grinder, the dried sheets were broken down into small ingestible particles and allowed to dry completely.

**Experimental Design.** Two independent experiments were conducted for 6% and 60% GA supplementation. For both experiments, fish were divided into two treatment groups (Control and GA) and maintained under Control or GA supplementation conditions for two weeks. For the 6% GA supplementation experiment, eight fish were housed in each 3.5 L tank at a 1:1 sex ratio (4 females and 4 males, total n = 142). For the 60% GA supplementation experiment, 16 fish were kept in an 8 L tank at a 1:1 sex ratio (8 females and 8 males, total n = 128). In both experiments, the Control fish were under the standard feeding regime of dry and live feed described earlier; whereas the GA fish were fed with dry feed supplemented with 6 or 60% GA and the live *Artemia*. To ensure proper nutrition and to avoid any caloric restrictions, all fish were fed *ad libitum* over 3% of their body weight/day (1). The dry feed was standardised by pre-weighing and then storing in labelled Eppendorf tubes as follows: 0.133 g for eight fish (6% GA) and 0.266 g for 16 fish (60% GA). The experiments were split into blocks where each block consisted of a Control tank and a GA tank (6% or 60% GA). We performed 9 blocks in the 6% GA experiment and 4 blocks in the 60% GA experiment (Fig. S1).

**Genomic DNA Extraction (gDNA) and 16S rRNA Sequencing.** Whole intestines were harvested from euthanised fish by dissection on ice and flash frozen in liquid nitrogen before storage at -80 °C. Before gDNA extraction, frozen samples were homogenized with a handheld homogeniser and pestle and kept on ice. Sequencing libraries were produced with 16S Amplicons V3-V4 by Source Bioscience, Cambridge, UK. The adapter sequences are listed in Table S1. The first ten samples from our first two blocks were sequenced on a MiSeq instrument with 250 base pair (bp) paired-end reads. A PhiX spike-in of 20% was used, and samples were sequenced on one lane on a MiSeq instrument. A further 40 samples were then prepared to produce sequence libraries with IDT for Illumina-Nextera DNA Unique Dual Indexes, using 16S Amplicons V3-V4 and then sequenced in separate sequencing experiment with 300 bp paired-end reads again on a MiSeq (762) instrument. Sequence quality was initially checked with FastQC (https://www.bioinformatics.babraham.ac.uk/projects/fastqc/). Using QIIME2, the sequences were trimmed, and adapters were removed using *cutadapt* in the QIIME2 environment. Samples were demultiplexed and *Dada2* was used to denoise and to trim using the following parameters "--p-trunc-len-f 240 \ --p-trunc-len-r 240 \--p-trim-left-f 13 \--p-trim-left-r 13 \". Feature tables were then generated and visualised before phylogenetic diversity analysis. Taxonomic assignment was performed using the SILVA database release 132 (https://www.arbsilva.de/fileadmin/silva_databases/qiime/Silva_132_release.zip) (2). Reads were delineated from the database based on the 341 and 806 primers listed in (Table S1). Our *silva* database was then trained using "qiime feature-classifier fit-classifier-naive-bayes". Taxonomy was assigned with qiime *sklearn* and data was exported from the QIIME2 environment to perform the downstream analysis using the package *phyloseq.*

**Metabolites Extraction and $^1$H NMR Metabolomic Analysis.** At the end of the 60% GA experiment, we collected intestines and brains from 16 females and 16 males by dissection on ice for further analysis (Table S2). Immediately after dissection, tissues were flash frozen in liquid nitrogen and stored at - 80 °C until further processing. In brief, the samples were weighed and placed in labelled Eppendorf tubes before adding some glass beads to them. Working on ice, 200 µL of ice-cold methanol and 42.5 µL of ultra-pure cold water were added to each tube before vortexing them. The samples were disrupted using a tissue lyser. After lysis, 100 µL of ice-cold chloroform were added and the tubes were vortexed again before adding another 100 µL of ice-cold chloroform in addition to 100 µL of ultra-pure cold water. The tubes were vortexed one more time and allowed to sit on ice for 15 min, followed by centrifugation at maximum speed for 5 min creating two visible layers. 250 µL of the top aqueous phase were transferred to new labelled Eppendorf tubes and left to dry gently in an oven for 48 hrs. The bottom chloroform layer was also collected into different new labelled tubes, evaporated, and stored. On the day of the NMR measurement, the aqueous layers were reconstituted in 550 µL of NMR phosphate buffer solution. All samples were then vortexed, and 500 µL was transferred to a 5 mm NMR tube for spectral acquisition.

**Phenotypic Characterisation and Behavioural Assessment (AI).** To initiate natural spawning, the fish were placed in pairs of opposite sex the day before in breeding tanks and kept separated by a divider. At 7:30 AM the next day, the divider was removed, and an hour of no disturbance was allocated to instigate the breeding in the zebrafish. Irrespective of the spawning outcome, all fish were placed back into their allocated system tanks by midday. The resulting clutches were assessed for various fitness traits including fertilisation success at 2 hpf, survival at 2 and 24 hpf, and the presence or absence of any developmental abnormalities (3) at 2 and 24 hpf. For the AI recording, the fish were placed in a round tank with a diameter of 235 mm and containing 2 L of system water. For the videos, a 40-minute clip was obtained by cropping out the first and last 10 minutes of each one-hour-long recording to reduce biases caused by the new environment and to standardise the footage. They were then further cropped for AI analysis. A total of 40 clips (5 clips x 2 treatments x 4 blocks), each lasting 5 min, were attained before each clip was separately analysed using the default idtracker.ai documentation.

**R analysis.** The following additional packages were used to generate plots, to handle data, and to perform statistical analysis: *ggplot* 2 3.4 (4), *Ape* 5.7-1 PMID: 30016406 (5), *Readxl* 1.4.3 (6), *dplyr* 1.1.2 (7), *Tibble* 3.2.1 (8), *Vegan* 2.6-4 (9), *ggThemeAssist* 0.1.5 (10), *ggpubr* 0.6.0.999 (11), *paletteer* 1.5.0 (12), *tidyverse* 2.0.0 (13), *pairwiseAdonis* 0.4.1 (14). The comprehensive R script used for statistical analysis and figure generation in this study can be provided upon request.

#### Supplementary figures

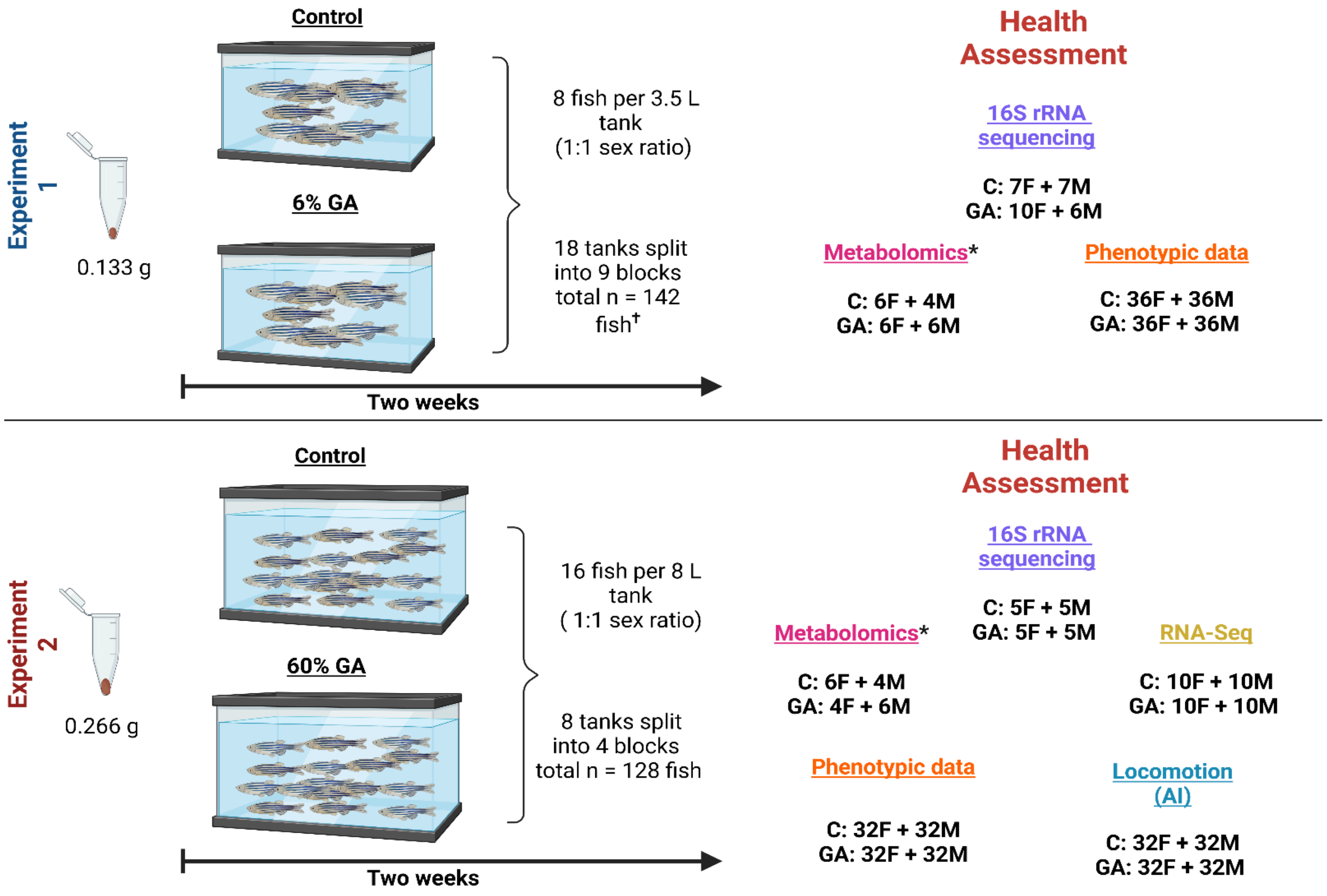


**Fig. S1.** Overview of the experimental design for both experiments. The total number and sex (F: female, M: male) of the fish used in the two independent experiments and the sample size for each health assessment. †Two fish died, *Controls were pooled from both experiments. The figure was created using BioRender.

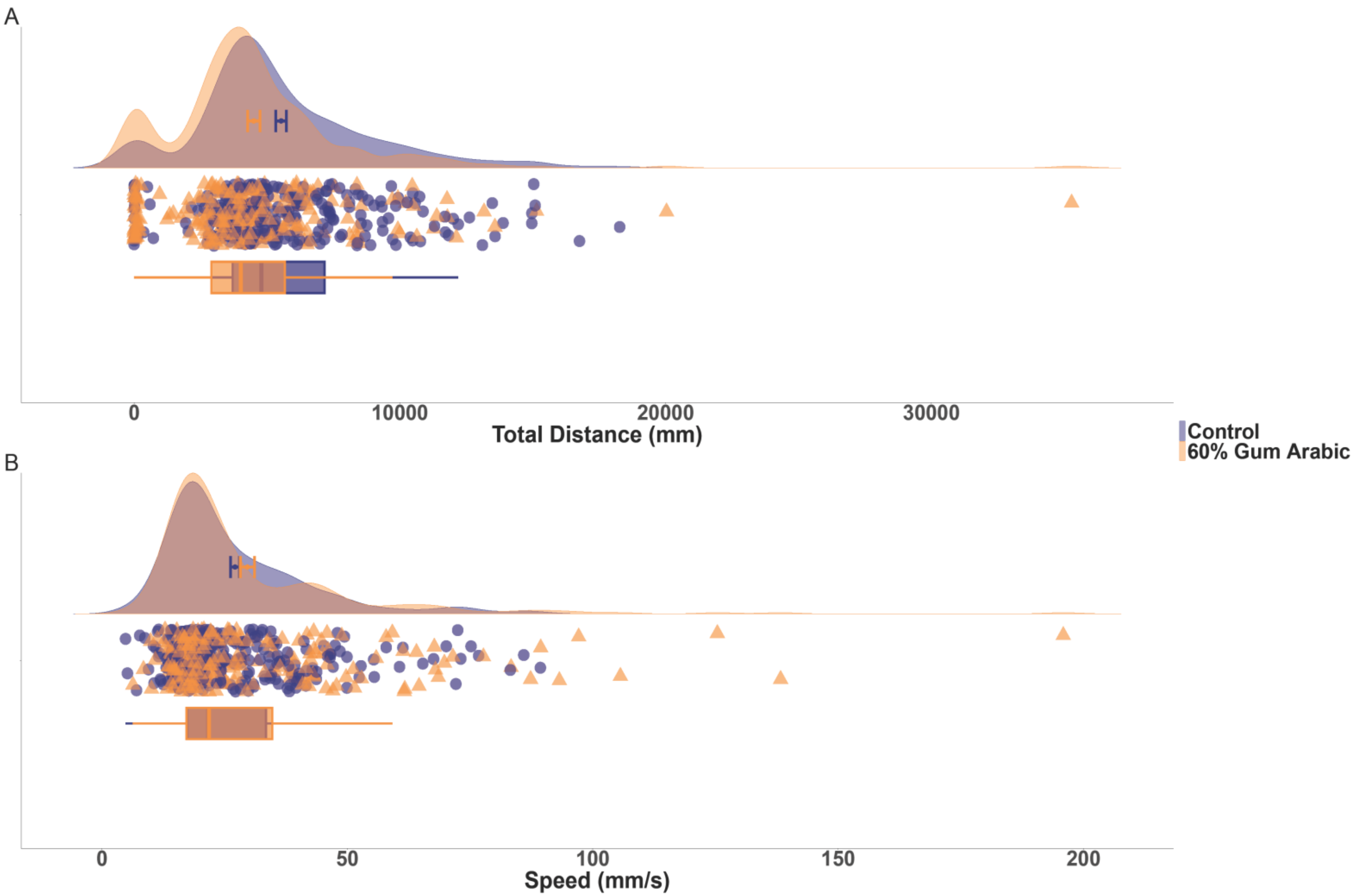


**Fig. S2.** The 60% GA fish swam faster yet covered a shorter distance compared to the Control fish. (*A* and *B*) Raindrop plots where each central point shows the mean value with the error bars representing the standard error of the mean (SEM) and each box plot represents the median, interquartile range, minimum, and maximum values of all the samples.

### Supplementary tables

**Table S1.** V3-V4 16S primer sequences and adapter used for library preparation and analyses.

| Primer | Sequence (5' -> 3') | Reference |
|---|---|---|
| 16s amplicon PCR primer forward | TCGTCGGCAGCGTCAGATGTGTATAAGAGACAGCCTACGGGNGGCWGCAG | (15) |
| 16s amplicon PCR primer reverse | GTCTCGTGGGCTCGGAGATGTGTATAAGAGACAGGACTACHVGGGTATCTAATCC | (15) |
| V3-V4 non overhang primer 341F | CCTACGGGNGGCWGCAG | |
| V3-V4 non overhang primer 806R | GACTACHVGGGTATCTAATCC | |
| Forward adapter | CTGTCTCTTATACACATCT | |
| Reverse adapter | CTGTCTCTTATACACATCT | |

**Table S2.** Supplemental information on the samples used in the metabolomics analysis.

| Pooled Sample | Individual Sample ID | Weight (mg) |
|---|---|---|
| Control Female Brain | 47.1 + 48.1 + C22.1 | 8.1 |
| Control Female Intestine | 47.5 + 48.5 + C22.5 | 23 |
| Control Male Brain | C23.1 + 75.1 | 5 |
| Control Male Intestine | C23.5 + 75.5 | 19.1 |
| 6% GA Female Brain | T33.1 + T35.1 + T39.1 | 10.8 |
| 6% GA Female Intestine | T33.5 + T35.5 + T39.5 | 62 |
| 6% GA Male Brain | 42.1 + 45.1 +T38.1 | 6.9 |
| 6% GA Male Intestine | 42.5 + 45.5 + T38.5 | 25 |
| 60% GA Female Brain | 110.1 + 91.1 | 3 |

| | | |
|---|---|---|
| 60% GA Female Intestine | 110.5 + 91.5 | 11.3 |
| 60% GA Male Brain | 84.1 + 90.1 + 106.1 | 4.1 |
| 60% GA Male Intestine | 84.5 + 90.5 + 106.5 | 12.3 |

**Table S3.** Supplemental information on the metabolites detected by $H^1$ NMR.

| Concentrations ($mmol.kg^{-1}$) in tissue specimen | BRAIN | BRAIN | BRAIN | BRAIN | BRAIN | BRAIN | INTESTINE | INTESTINE | INTESTINE | INTESTINE | INTESTINE | INTESTINE |
|---|---|---|---|---|---|---|---|---|---|---|---|---|
| Sex | M | F | M | F | M | F | M | F | M | F | M | F |
| Diet | C | C | 6% | 6% | 60% | 60% | C | C | 6% | 6% | 60% | 60% |
| 2-Oxoglutarate | 0.14 | 0.56 | 0.62 | 0.48 | 0.66 | 0.55 | 0.00 | 0.00 | 0.00 | 0.00 | 0.00 | 0.00 |
| 4-Aminobutyrate | 0.64 | 2.45 | 1.88 | 1.48 | 1.87 | 2.29 | 0.17 | 0.32 | 0.19 | 0.03 | 0.00 | 0.45 |
| adenosine phosphate | 0.07 | 0.32 | 0.28 | 0.36 | 0.00 | 0.04 | 0.00 | 0.00 | 0.00 | 0.00 | 0.00 | 0.00 |
| Acetate | 1.79 | 5.49 | 6.14 | 4.01 | 9.39 | 12.89 | 1.81 | 1.84 | 1.83 | 0.62 | 2.37 | 3.23 |
| Adenine | 0.11 | 0.31 | 0.38 | 0.46 | 0.84 | 0.79 | 0.28 | 0.30 | 0.21 | 0.06 | 0.00 | 0.36 |
| Alanine | 0.16 | 0.80 | 0.72 | 0.56 | 0.83 | 1.19 | 5.27 | 6.59 | 4.10 | 1.07 | 0.86 | 7.09 |
| Arginine | 0.00 | 0.00 | 0.00 | 0.00 | 0.00 | 0.00 | 2.65 | 1.51 | 2.20 | 0.57 | 0.51 | 3.92 |
| Asparagine | 0.00 | 0.00 | 0.00 | 0.00 | 0.00 | 0.00 | 1.55 | 1.90 | 1.25 | 0.41 | 0.43 | 2.22 |
| Aspartate | 0.13 | 0.70 | 1.06 | 0.70 | 0.99 | 0.99 | 1.57 | 1.79 | 1.19 | 0.44 | 0.43 | 1.56 |
| N-acetylaspartylglutamate | 4.81 | 20.32 | 17.44 | 12.89 | 28.71 | 22.20 | 0.00 | 0.00 | 0.00 | 0.00 | 0.00 | 0.00 |
| Betaine | 0.13 | 0.42 | 0.44 | 0.49 | 0.65 | 0.77 | 0.36 | 0.34 | 0.15 | 0.03 | 0.05 | 0.52 |
| Choline | 0.04 | 0.21 | 0.20 | 0.17 | 0.25 | 0.46 | 0.71 | 0.40 | 0.39 | 0.11 | 0.00 | 0.74 |
| Citrulline | 0.00 | 0.00 | 0.00 | 0.00 | 0.00 | 0.00 | 0.47 | 0.65 | 0.58 | 0.35 | 0.00 | 1.51 |
| Creatine | 1.65 | 6.39 | 6.92 | 6.23 | 7.30 | 9.72 | 2.17 | 2.90 | 2.53 | 0.71 | 0.18 | 2.92 |
| Dimethylamine | 0.03 | 0.23 | 0.13 | 0.25 | 0.19 | 0.02 | 0.25 | 0.24 | 0.30 | 0.15 | 0.00 | 0.50 |

| | | | | | | | | | | | | |
|---|---|---|---|---|---|---|---|---|---|---|---|---|
| Ethanol | 0.14 | 0.39 | 0.36 | 0.27 | 0.66 | 0.64 | 0.00 | 0.00 | 0.00 | 0.00 | 0.00 | 0.00 |
| Formate | 0.98 | 2.41 | 3.32 | 2.08 | 6.35 | 7.17 | 1.12 | 1.07 | 0.78 | 0.33 | 2.04 | 1.90 |
| Fumarate | 0.00 | 0.00 | 0.00 | 0.00 | 0.00 | 0.00 | 0.05 | 0.06 | 0.03 | 0.02 | 0.00 | 0.07 |
| Glucose | 0.00 | 0.00 | 0.00 | 0.00 | 0.00 | 0.00 | 14.30 | 15.59 | 6.10 | 1.53 | 1.05 | 7.73 |
| Glutamate | 0.59 | 2.06 | 2.29 | 1.79 | 2.79 | 2.33 | 2.80 | 3.56 | 2.91 | 0.78 | 0.14 | 4.01 |
| Glutamine | 0.36 | 1.46 | 1.66 | 1.69 | 2.15 | 2.11 | 2.85 | 2.42 | 2.17 | 0.67 | 0.39 | 4.16 |
| Glutathione | 0.00 | 0.00 | 0.00 | 0.00 | 0.00 | 0.00 | 0.26 | 0.22 | 0.24 | 0.05 | 0.00 | 0.31 |
| Glycerol | 0.35 | 1.14 | 1.30 | 1.09 | 1.65 | 12.36 | 1.83 | 2.99 | 2.10 | 0.60 | 0.76 | 1.37 |
| Glycine | 0.19 | 1.06 | 1.01 | 0.73 | 1.16 | 1.43 | 1.96 | 2.33 | 1.36 | 0.59 | 0.55 | 2.59 |
| Guanosine | 0.00 | 0.00 | 0.00 | 0.00 | 0.00 | 0.00 | 0.18 | 0.18 | 0.12 | 0.03 | 0.00 | 0.15 |
| Histidine | 0.30 | 0.76 | 1.43 | 0.64 | 1.47 | 2.51 | 0.85 | 1.11 | 0.64 | 0.17 | 0.00 | 0.41 |
| Inosine phosphate | 0.00 | 0.00 | 0.00 | 0.00 | 0.00 | 0.00 | 0.11 | 0.10 | 0.13 | 0.10 | 0.00 | 0.13 |
| Inosine | 0.11 | 0.21 | 0.31 | 0.36 | 0.34 | 0.95 | 1.23 | 1.13 | 0.64 | 0.20 | 0.21 | 1.09 |
| Isoleucine | 0.03 | 0.06 | 0.09 | 0.12 | 0.11 | 0.26 | 2.34 | 2.86 | 1.80 | 0.44 | 0.39 | 2.93 |
| Lactate | 1.62 | 6.75 | 11.22 | 6.74 | 8.65 | 11.04 | 1.97 | 2.09 | 1.67 | 0.45 | 0.28 | 2.54 |
| Leucine | 0.03 | 0.06 | 0.09 | 0.15 | 0.18 | 0.37 | 4.92 | 6.84 | 4.52 | 1.10 | 0.82 | 7.39 |
| Lysine | 0.00 | 0.00 | 0.00 | 0.00 | 0.00 | 0.00 | 5.95 | 6.94 | 4.05 | 0.85 | 0.33 | 7.67 |
| Maltose | 0.00 | 0.00 | 0.00 | 0.00 | 0.00 | 0.00 | 0.98 | 0.34 | 0.00 | 0.09 | 0.00 | 0.00 |
| Methanol | 0.21 | 0.60 | 0.68 | 0.40 | 1.18 | 1.61 | 0.29 | 0.23 | 0.16 | 0.06 | 0.29 | 0.37 |
| Methionine | 0.03 | 0.09 | 0.09 | 0.07 | 0.17 | 0.11 | 1.62 | 2.21 | 1.45 | 0.29 | 0.17 | 2.34 |
| N-Acetylcysteine | 0.00 | 0.00 | 0.00 | 0.00 | 0.00 | 0.00 | 0.36 | 0.21 | 0.22 | 0.08 | 0.00 | 0.32 |
| Niacinamide | 0.00 | 0.00 | 0.00 | 0.00 | 0.00 | 0.00 | 0.15 | 0.21 | 0.11 | 0.11 | 0.00 | 0.31 |
| N-Acetylaspartate | 0.55 | 3.06 | 2.92 | 2.15 | 3.09 | 3.10 | 0.00 | 0.00 | 0.00 | 0.00 | 0.00 | 0.00 |
| O-Acetylcarnitine | 0.01 | 0.03 | 0.04 | 0.02 | 0.04 | 0.06 | 0.02 | 0.01 | 0.08 | 0.02 | 0.00 | 0.04 |
| O-Acetylcholine | 0.01 | 0.03 | 0.04 | 0.02 | 0.04 | 0.06 | 0.00 | 0.00 | 0.00 | 0.00 | 0.00 | 0.00 |
| O-Phosphocholine | 0.03 | 0.14 | 0.13 | 0.11 | 0.19 | 0.18 | 2.21 | 0.79 | 0.86 | 0.27 | 0.16 | 1.88 |
| Ornithine | 0.00 | 0.00 | 0.00 | 0.00 | 0.00 | 0.00 | 0.07 | 0.30 | 0.41 | 0.04 | 0.00 | 0.20 |
| Phenylalanine | 0.00 | 0.00 | 0.00 | 0.00 | 0.00 | 0.00 | 2.72 | 3.07 | 1.67 | 0.37 | 0.28 | 2.43 |
| Proline | 0.00 | 0.00 | 0.00 | 0.00 | 0.00 | 0.00 | 1.67 | 2.07 | 1.54 | 0.44 | 0.50 | 2.51 |
| Serine | 0.10 | 0.27 | 0.24 | 0.33 | 0.34 | 0.40 | 2.98 | 3.73 | 3.35 | 0.92 | 0.22 | 4.92 |
| Succinate | 0.01 | 0.01 | 0.03 | 0.05 | 0.07 | 0.07 | 0.05 | 0.08 | 0.06 | 0.01 | 0.00 | 0.05 |
| Sucrose | 0.00 | 0.00 | 0.00 | 0.00 | 0.00 | 0.00 | 0.35 | 0.00 | 0.00 | 0.00 | 0.00 | 0.00 |
| Taurine | 1.33 | 6.28 | 6.61 | 6.18 | 6.05 | 9.09 | 10.24 | 8.28 | 9.09 | 2.09 | 1.33 | 8.57 |
| Threonine | 0.00 | 0.00 | 0.00 | 0.00 | 0.00 | 0.00 | 1.68 | 2.08 | 1.81 | 0.52 | 0.20 | 2.97 |
| Tryptophan | 0.00 | 0.00 | 0.00 | 0.00 | 0.00 | 0.00 | 0.47 | 0.64 | 0.37 | 0.10 | 0.00 | 0.53 |
| Tyrosine | 0.00 | 0.00 | 0.00 | 0.00 | 0.00 | 0.00 | 2.70 | 3.06 | 2.16 | 0.41 | 0.41 | 4.46 |

| | | | | | | | | | | | | |
|---|---|---|---|---|---|---|---|---|---|---|---|---|
| UDP-galactose | 0.00 | 0.00 | 0.00 | 0.00 | 0.00 | 0.00 | 0.37 | 0.19 | 0.24 | 0.07 | 0.00 | 0.25 |
| UDP-glucose | 0.00 | 0.00 | 0.00 | 0.00 | 0.00 | 0.00 | 0.79 | 0.33 | 0.39 | 0.14 | 0.00 | 0.50 |
| Uracil | 0.00 | 0.00 | 0.00 | 0.00 | 0.00 | 0.00 | 0.32 | 0.17 | 0.24 | 0.05 | 0.00 | 0.50 |
| Uridine | 0.00 | 0.00 | 0.00 | 0.00 | 0.00 | 0.00 | 0.11 | 0.12 | 0.09 | 0.02 | 0.00 | 0.19 |
| Valine | 0.03 | 0.09 | 0.07 | 0.12 | 0.19 | 0.28 | 3.21 | 3.75 | 2.48 | 0.67 | 0.44 | 4.24 |
| Xanthine | 0.64 | 0.34 | 2.87 | 0.94 | 4.11 | 3.69 | 0.00 | 0.17 | 0.16 | 0.35 | 0.66 | 0.24 |
| Citrate | 0.03 | 0.14 | 0.17 | 0.18 | 0.11 | 0.24 | 0.13 | 0.10 | 0.11 | 0.02 | 0.00 | 0.17 |
| Malate | 0.11 | 0.31 | 0.50 | 0.42 | 1.06 | 1.03 | 0.82 | 0.51 | 0.48 | 0.15 | 0.00 | 0.73 |
| myo-Inositol | 0.20 | 1.05 | 0.97 | 0.76 | 1.38 | 1.05 | 0.00 | 0.00 | 0.00 | 0.00 | 0.00 | 0.00 |
| sn-Glycero-3-phosphocholine | 0.07 | 0.28 | 0.50 | 0.31 | 0.56 | 0.66 | 1.09 | 4.28 | 1.77 | 0.19 | 0.00 | 2.32 |

**Table S4.** Supplemental information on the statistical models.

| | **Clutch production (6% GA)** | | | | **Clutch production (60% GA)** | | | |
|---|---|---|---|---|---|---|---|---|
| **Effect** | **Est** | $x^2$ | **DF** | $p$ | **Est** | $x^2$ | **DF** | $p$ |
| GA treatment | 0.36 | 0.40 | 1 | 0.52 | -0.41 | 0.54 | 1 | 0.46 |
| Sex | 0.31 | 0.35 | 1 | 0.55 | -0.27 | 0.27 | 1 | 0.61 |
| Treatment: Sex | -0.49 | 0.47 | 1 | 0.49 | 1.32 | 3.18 | 1 | 0.075 |
| | **Random effects variance: Tank: 0.23, SD=0.48** | | | | **Random effects variance: Tank: 0.05, SD= 0.24** | | | |

| | **Total egg number (6% GA)** | | | | **Total egg number (60% GA)** | | | |
|---|---|---|---|---|---|---|---|---|
| **Effect** | **Est** | $x^2$ | **DF** | $p$ | **Est** | $x^2$ | **DF** | $p$ |
| GA treatment | -0.19 | 0.10 | 1 | 0.76 | 0.34 | 0.13 | 1 | 0.72 |
| Sex | -0.16 | 0.02 | 1 | 0.88 | 0.26 | 0.75 | 1 | 0.39 |
| Treatment: Sex | 0.22 | 0.22 | 1 | 0.64 | 0.37 | 12.56 | 1 | < 0.001 *** |
| | **Random effects variance:<br>Observation: 0.74, SD = 0.86<br>Tank: <0.001, SD = <0.001<br>2h counter: 0.03, SD = 0.16** | | | | **Random effects variance:<br>Observation: 0.35, SD = 0.60<br>Tank: 0.06, SD = 0.24<br>2h counter: 0.15, SD = 0.39** | | | |

| | **Unfertilised embryos (6% GA)** | | | | **Unfertilised embryos (60% GA)** | | | |
|---|---|---|---|---|---|---|---|---|
| **Effect** | **Est** | ***$x^2$*** | **DF** | ***p*** | **Est** | ***$x^2$*** | **DF** | ***p*** |
| GA treatment | -0.27 | 0.08 | 1 | 0.77 | 1.23 | 0.39 | 1 | 0.53 |
| Sex | 0.19 | 0.26 | 1 | 0.61 | -0.14 | 4.57 | 1 | 0.03 * |
| Treatment: Sex | 0.21 | 0.03 | 1 | 0.86 | -1.77 | 2.92 | 1 | 0.09 |
| | **Random effects variance:<br>ID: 3.3, SD = 1.82<br>2h counter: 0.21, SD = 0.46** | | | | **Random effects variance:<br>ID: 1.49, SD = 1.2<br>2h counter: 1.15, SD = 1.7** | | | |

| | **Dead embryos 24h (6% GA)** | | | | **Dead embryos 24h (60% GA)** | | | |
|---|---|---|---|---|---|---|---|---|
| **Effect** | **Est** | ***$x^2$*** | **DF** | ***p*** | **Est** | ***$x^2$*** | **DF** | ***p*** |
| GA treatment | -0.88 | 0.55 | 1 | 0.46 | 0.64 | 0.82 | 1 | 0.37 |
| Sex | -1.02 | 1.54 | 1 | 0.22 | 0.57 | 1.94 | 1 | 0.16 |
| Treatment: Sex | 1.02 | 0.1.77 | 1 | 0.18 | 0.80 | 6.31 | 1 | 0.01 * |
| | **Random effects variance:<br>Observation: 1.6, SD = 1.3<br>Tank: 0.23, SD = 0.5** | | | | **Random effects variance:<br>Observation: 1.5, SD = 1.2<br>Tank: 0.06, SD = 0.2<br>2h Counter: 0.10, SD = 0.33** | | | |
